# Biometric aging clock for practical utility in clinical settings

**DOI:** 10.1101/2025.04.14.648594

**Authors:** Deepankar Nayak, Fedor Galkin, Subir Kumar Ghosh

## Abstract

Aging clocks are an essential tool for biogerontological research that enables a quick assessment of one’s pace of aging. In research settings, clocks trained on -omics data types are the most popular since they allow for an inspection of the most basic cellular processes of aging. In clinical settings, however, -omics biomarkers of aging are impractical since they are linked to extra logistical load and costs, and they require personnel training. In this article, we present a cost-efficient aging clock that can be implemented easily in most clinics and hospitals to measure patients’ aging rates. The clock requires only 22 biomarkers, including 17 blood test parameters, and four biometric measures (blood pressure, body mass index, and waist circumference). The clock predicts one’s chronological age with a mean average error of 7.35 years, and it reveals associations with hypertension, cancer, and obesity.

## Introduction

Biogerontology is a rapidly developing field of biology directed toward the discovery of interventions to prolong the human lifespan. The population of developed countries is rapidly aging, a phenomenon commonly recognized as a major economic and social concern. The global median age increased from 20 years in 1970 to 30 in 2022, with some countries—such as Japan—displaying extreme rates of population aging.^1^ This upward aging trend is coupled with an increased incidence of dangerous age-related health conditions, such as heart diseases, cancer, cirrhosis, and diabetes.^2^ To prevent the negative consequences of this process, numerous research teams and commercial organizations are developing geroprotectors—drugs and therapies that can prevent or reverse systemic damage accumulated during aging.

Compounds that can destroy senescent cells (senolytics) or rejuvenate them (senomorphics) are being discovered in screenings assisted by artificial intelligence (AI).^3,4^ Yet, even the most promising compounds identified in in silico experiments require proper validation during preclinical and clinical trials. Experiments on short-living model organisms, such as *C*.*elegans* and mice, commonly use the extension of average or maximal lifespans as an endpoint. Prospective experiments with animals contribute to the knowledge of geroprotectors’ mechanism of action, but their methodology cannot be translated into human trials.^5^ Since measuring a compound’s effect on a human lifespan is not feasible, developing a proxy for one’s longevity potential that may be obtained in a reasonable observation period becomes necessary.

Such proxies have been implemented via machine learning using various biodata types: transcriptomics, DNA methylation profiles, proteomics, and metagenomics.^6–10^ These statistical models of aging, or aging clocks, enable researchers to study the molecular basics of the aging process. However, they prove hard to adopt in clinical practice since they require complex sample-processing techniques and expensive equipment, such as sequencing machines. Clinics aiming to adopt the latest longevity medicine developments must, therefore, invest in new lab facilities and training. Alternatively, such clinics may choose to partner with a specialized service provider who would manage -omics data generation. This approach is paired with managing collecting patient consent, data transfer agreements, and extra administration efforts due to increased complexity in a clinic’s logistics. From this point of view, aging clocks that can interpret the biodata types native to the clinical practice are an optimal solution for both price-conscious patients and professionals seeking operational convenience.

Previously, we released aging clocks that can interpret clinical blood tests in the aging context ^11^. The pace of aging detected with these clocks is associated with mortality risk, psychological well-being, and aging-related diseases.^12,13^ While these blood aging clocks rely only on standard clinical tests, the number of features used in the models still significantly limits their scope for practitioners. A patient’s willingness to pay (WTP) for diagnostic procedures remains low, with a median WTP below 100 USD even for such procedures as HIV testing and breast or colon cancer screening.^14^ Hence, lowering the cost and number of parameters in a pace-of-aging blood panel is essential for patients’ adoption of aging clocks.

In this article, we present a new blood aging clock that requires only **17** blood parameters compatible with blood panels prescribed in clinical practice (**Figure 1**). We significantly reduced the feature number by including biometric measures such as blood pressure, body mass index (BMI), and waist circumference, which may be measured at no additional cost. The aging clock reported in this article may be used to affordably measure the pace of aging, and it fits better with retrospective studies in which additional data collection is not feasible.

**Figure 1.**
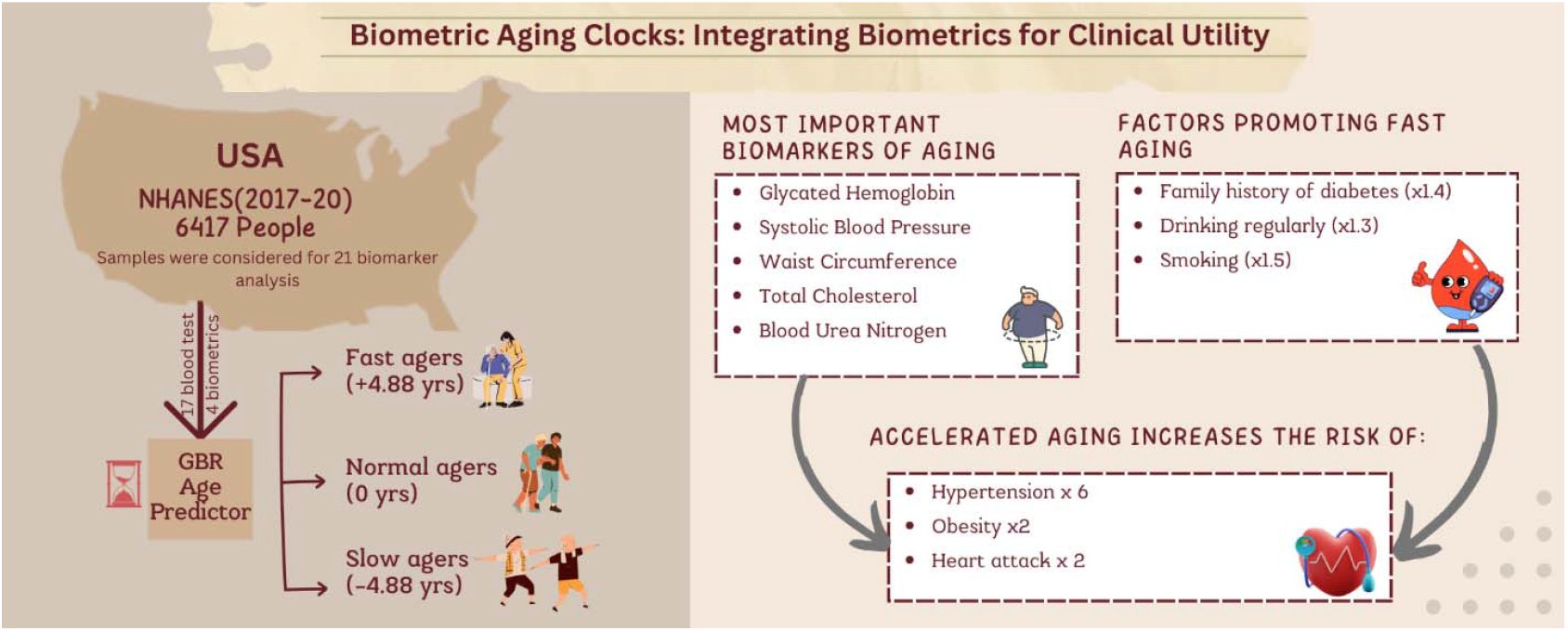
This study presents a new biometric aging clock developed with data from NHANES. This aging clock is based on 21 clinically relevant features

## Methods

### Study cohort

For this study, we used publicly available data from the National Health and Nutrition Examination Survey (NHANES) 2017–2020 waves, containing 15,560 individual records in pre-COVID-19 pandemic files. After the participants who missed physical examinations, were pregnant, or were outside the 20– 65-year age range were removed, 6,417 samples were considered for further analysis. Among them, 5,542 samples were defined as “healthy” and 875 as “unhealthy,” based on the presence of any of the following health conditions in the original annotation: heart stroke, heart attack, coronary heart disease, congestive heart failure, liver diseases, or cancer malignancy.

According to the NHANES guidelines, liver diseases include viral hepatitis (including hepatitis A, hepatitis B, and hepatitis C), autoimmune liver disease (including primary biliary cirrhosis, autoimmune hepatitis, and sclerosing cholangitis), genetic liver diseases (including alpha-1-antitrysin deficiency, hemochromatosis, and Wilson’s disease), drug- or medication-induced liver disease, alcoholic liver disease, non-alcoholic fatty liver disease, fatty liver disease, liver cancer, liver cyst, liver abscess, liver fibrosis, and liver cirrhosis. The healthy cohort was split into the training set (*n* = *4156, 75%*) and the test set (*n* = *1,386, 25%*). The unhealthy cohort was later used as a discovery set.

### Feature selection

The NHANES 2017–2020 laboratory and physical examination tables contain 124 variables describing blood biomarkers and biometric measurements. Repeated measurements (such as blood pressure or heart rate) were replaced with mean values across all observations, after which we excluded variables with >10% missing values to obtain a list of 55 variables. We then removed the collinear variables based on Pearson’s r cutoff (*r < 0*.*95*) for pairwise correlation. To consider only the variables containing aging-related information, chronological age was regressed as a linear function of single features, and all features with insignificant (*P-value > 0*.*25*) coefficients were discarded. Finally, we removed the features based on a variance inflation factor cutoff (*VIF > 10*) to obtain a list of 44 variables that were considered for model training.

### Model Training

To train the aging clock, we used the GradientBoostingRegressor (GBR) module from the sklearn library for Python. A preliminary model was trained on 44 features that remained after filtering to predict the age of the NHANES participants. The preliminary model’s parameters were set using a random grid search. After the impact of each feature on the prediction with permutation feature importance (PFI) was assessed, the least important features whose aggregate importance composed < 10% of the observed total importance were removed, yielding a list of 25 features.

To finalize the model, we trained GBR using the <u>Hyperopt</u> Python library to fine-tune the parameters of each iteration, including the number of features (varying their number from the 20^th^ to 27^th^ most important ones). The best-performing iteration was determined based on the mean absolute error (MAE) achieved in five-fold cross-validation. Thus, the final model reported in this article was trained with the following parameters: alpha = 0.800, learning_rate = 0.060, loss = “huber,” max_depth = 3, max_features = “sqrt,” min_samples_split = 50, criterion = “friedman_mse,” validation_fraction = 0.1, n_iter_no_change = 5, warm_start = True, n_estimators = 450, and subsample = 0.748.

The performance of the GBR model was compared against an elastic net (EN) model trained on the same split with the same variables. The EN model was implemented using the <u>ElasticNet</u> module from the sklearn library for Python.

### Statistical tests

All model performance metrics were measured with the sklearn.metrics module for Python. For the comparison of averages, the Mann–Whitney U-test from scipy.stats was used.

To enable comparisons of age prediction between different age groups, we adjusted each prediction by deducting the average prediction error for each observation’scorresponding five-year age group. To calculate odds ratios (ORs), we defined people with an adjusted prediction error above +*4*.*88* years as “fast agers” and those with an error below −*4*.*88* years as “slow agers.” The threshold error was selected as the MAE of adjusted predictions in the training set.

The ORs and corresponding confidence intervals (CIs) were derived from a custom implementation of McNemar’s paired test. Fast agers from the test set were paired with age-and-sex-matched counterparts among slow agers.

## Results

### Biometric clock accurately predicts chronological age

We trained the combined blood and biometric aging clock using a collection of 4,156 NHANES entries, obtaining a cross-validation accuracy of *MAE* = 7.09 years and 7.70 years in a test set comprising 1,386 samples of participants who were considered healthy (**Table 1 and Figure 2**). The model, however, underperformed in the discovery set, which contained only participants with serious health disorders (n = 875)—*MAE* = 8.43 years. To measure the GBR model’s accuracy without sample effects, we applied the observation weights specified in NHANES to the prediction errors in the test cohort, and we obtained an MAE of 7.35 years. Individual-level predictions are available in **Supplementary File 1**.

**Table 1.** The performance of the GBR age predictor with the training and validation cohorts compared to the baseline (mean age assignment) and an elastic net model. The weighted MAEs were calculated based on the prediction errors weighted according to the sample weights provided by NHANES. EN — Elastic Net, GBR — Gradient Boosting Regressor, MAE — Mean Absolute Error, R^2^ —coefficient of determination, Q1 — first quartile value, Q3 — third quartile value

|  |  | Training set | Test set | Discovery set |
| --- | --- | --- | --- | --- |
| GBR | $R^2$ | 0.55 | 0.48 | < 0 |
|  | MAE, years (weighted) | 7.09 (6.27) | 7.70 (7.35) | 8.43 (8.31) |
| EN | $R^2$ | 0.37 | 0.38 | < 0 |
|  | MAE, years | 8.65 | 8.68 | 9.43 |
|  | Baseline MAE, years | 11.48 | 11.55 | 8.04 |
|  | n, participants | 4,156 | 1,386 | 875 |
|  | Median age (Q1;Q3), years | 42 (31;55) | 42 (31;54) | 56 (49;61) |

**Figure 2.**
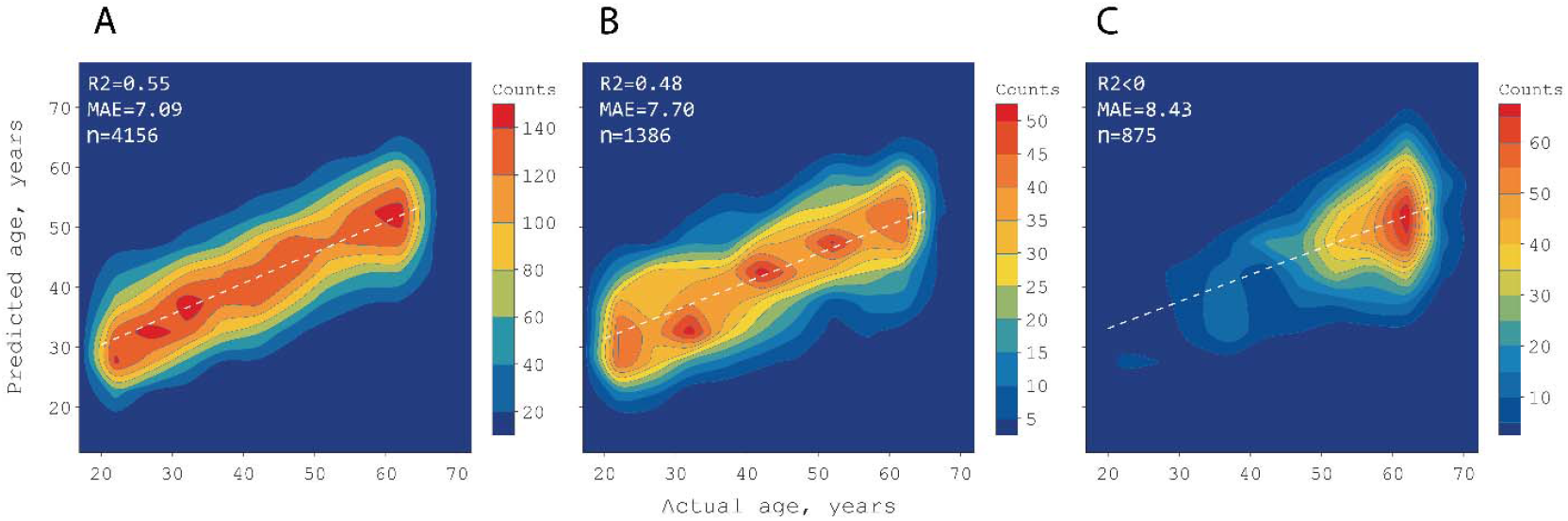
Density map representing the performance of the GBR age predictor in (A) cross-validation, (B) the test set comprising participants without major health conditions, and (C) participants with a history of heart disease, liver disease, or malignancy. The color bars indicate continuous areas containing the same number of people. “MAE” stands for mean absolute error, and “R^2^”stands for the coefficient of determination.

The trained model showed a significant improvement over a simpler EN implementation based on the same variables (*MAE = 8*.*68* for the test set).

The predictions for male participants were, on average, 0.84 years higher than those for the female participants in the test set (*P-value < 0*.*05*). Conversely, different age groups displayed significant (*P-value < 0*.*001*) deviations in their error distributions. While the youngest participants’ (20–25 years) predictions were, on average, 10.01 years above their actual ages, the oldest participants (60–65 years) had an average prediction error of −11.40 years. Hence, we implemented an age-group-based adjustment to enable a comparison of model predictions between different age groups in the following sections.

### Relative importance of aging biomarkers

We used the PFI technique to see which parameters the model considered the most important for age prediction (**Figure 3**). Among the top five most important features, we found two biometric markers (systolic blood pressure and waist circumference) and three blood biomarkers (glycated hemoglobin, total cholesterol, and blood urea nitrogen, BUN). The most influential biomarker was glycated hemoglobin; its permutation resulted in an MAE increase of 1.47 years.

**Figure 3.**
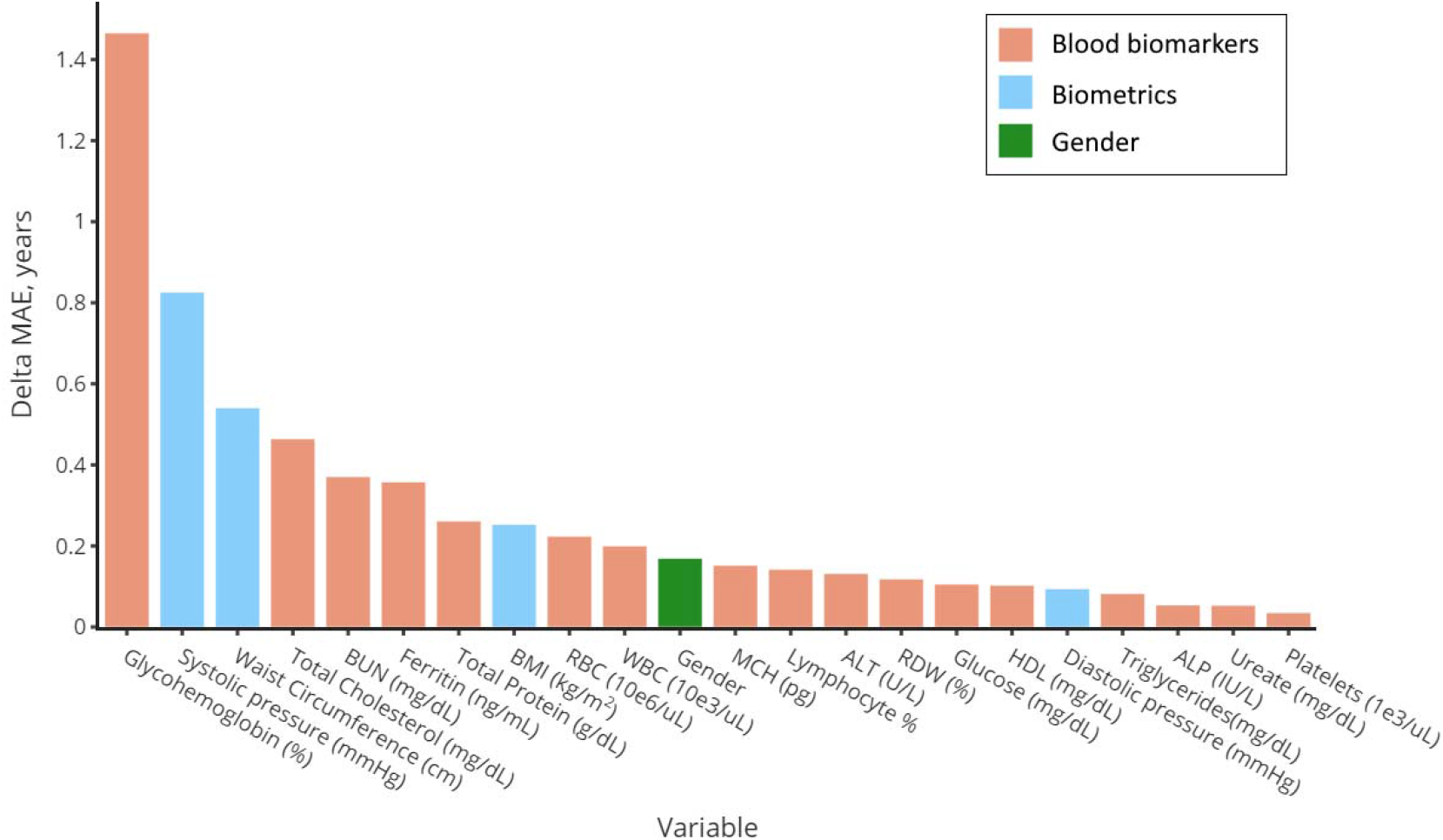
The results of the PFI analysis: glycated hemoglobin was the most influential variable. Its permutation decreased the model’s performance by 1.47 years. MAE — Mean Absolute Error

### Comorbidities are associated with an accelerated aging rate

To confirm whether the trained aging clock’s predictions conveyed information about a person’s health status, we sought statistical associations between the prediction error and certain diagnoses.

We paired fast agers (prediction error > 4.88 years) from the test set with age-and-sex-matched slow agers (prediction error < −4.88 years), and we applied McNemar’s test to obtain the ORs. We identified the increased pace of aging as a factor predisposing individuals to hypertension (*OR = 5*.*91*) and obesity (*OR = 2*.*17*). No significant association was identified between the aging rate and thyroid disorders or gallstones (see Table 2).

**Table 2.** OR of having a certain condition if a participant was a fast ager (compared to age-and-sex-matched slow agers). *—*p-value* < 0.050; **—*p-value* < 0.010; ***—*p-value* < 0.001; “support”— number of paired observations. OR — Odds Ratio

| Condition | OR (95% CI) | Support |
| --- | --- | --- |
| Hypertension*** | 5.909 (3.742–9.755) | 334 |
| Obesity*** | 2.171 (1.483–3.224) | 290 |
| Heart attack** | 2.143 (1.347–3.486) | 571 |

To enable a similar assessment for life-threatening diseases, we applied McNemar’s test to the combined discovery and test sets. Of health conditions such as coronary heart disease, liver diseases, cancer, and heart attack, only heart attack showed a significant association with an increased pace of aging (*OR* = *2*.*14*; Table 2).

We further explored whether an individual’s familial medical history is associated with fast aging. Due to a low number of observations, we repeated McNemar’s test using the combined training and test sets. The presence of close relatives who had experienced a heart attack or suffered from asthma was not significantly associated with a participant’s pace of aging. However, participants with a family history of diabetes were 1.43 times more likely to be fast agers. Similarly, we established that drinking more than once a week (regular drinking) was also linked to accelerated aging, although total abstinence did not affect the odds of being a fast ager (*P-value > 0*.*05*). Occasionally or regularly smoking tobacco was also positively associated with fast aging (Table 3).

**Table 3.**
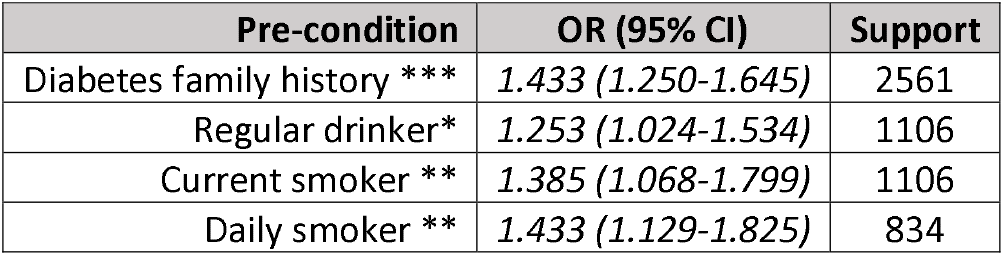
OR of being a fast ager if a participant displayed a pre-condition (compared to age-and-sex-matched slow and normal agers). *—*p-value* < 0.050; **—*p-value* < 0.010; ***—*p-value* < 0.001; “support”—number of paired observations. OR — Odds Ratio

## Discussion

In this article, we have developed and validated a novel aging clock using a combination of biometric and blood measurements. The model’s displayed accuracy (*MAE = 7*.*70*) is on par with that of similar models.^15^ Using the participants’ weight coefficients, as recommended by the NHANES maintainers, the presented aging clock is expected to have an *MAE of 7*.*35* years if applied to the scope of the US population. This performance, however, is likely to vary when applied to other populations. Previous studies of blood biomarkers have shown that an aging clock’s accuracy may decrease by > 50% when the tool is transferred to an out-of-scope population.^12^ This decrease may be due to transnational differences in public health policies, ethnicity-specific biochemical polymorphism, and the national regulation of clinical testing. Some studies have even suggested that general tests may be subject to major inter-individual baseline variability, hindering their interpretation.^16^ Additionally, liver biomarkers that are commonly used to indicate liver damage have reportedly shown inconsistent results. In 30% of observations showing abnormal alanine transaminase (ALT), aspartate transaminase (AST), and bilirubin levels, repeated measurements reclassify patients as having normal levels.^17^ The variability in singular blood biomarkers suggests that optimal performance demands repeated measurements in a short timeframe. However, any extra blood testing accompanies increased costs for patients, defeating the purpose of developing an affordable pace-of-aging biomarker panel. While intra-individual variability poses a challenge in a single-test setting, the most effective measure to counteract this difficulty lies in informing patients of which factors might affect their blood biomarkers immediately before blood drawing, such as sleep quality the night before, strenuous exercise, fatty food, alcohol intake, and the reproductive cycle ^18–21^.

While the blood parameters selected for this study’s aging clock are highly reliable in assessing the accumulated burden of aging-related damage to organisms, assessments of their utility as aging biomarkers must be regularly updated and reviewed. Even the more reproducible biomarkers have shifting generational distributions. In particular, glycated hemoglobin displayed a decreasing trend in the United Kingdom during the 1980–1990s across all age groups because a nationwide shift in food behaviors, lifestyle choices, and healthcare policies improved glycemic control.^22^ More recently, governments’ efficiency in implementing health policies, household incomes, and diabetes awareness have been suggested as factors affecting glycated hemoglobin differences across countries.^23^ Arguably, the same confounders evolving within one country due to economic and political factors could cause significantly different biomarker baselines across generations. The currently presented study was based on the latest NHANES wave; therefore, it should represent the current US population’s health. However, exploring whether including a year-of-birth (rather than chronological age) adjustment would yield more accurate age predictions and risk estimations would be worthwhile.

Biometric measures also display generational trends. In the United States, the portion of people with systolic blood pressure > 140 mmHg decreased from 37% to 33% between 2007 and 2016, supposedly due to increased antihypertensive prescriptions.^24^ Similarly, other cardiovascular health measures have trended downward; mean total cholesterol fell from 203 mg/dL in 2000 to 188 mg/dL in 2018, and one contributing factor was a higher rate of cholesterol screening.^25^ The inclusion of easy-to-obtain biometrics has allowed us to reduce aging rate assessment procedures’ invasiveness. While age prediction accuracy may increase with more blood-derived features, such an approach is also associated with higher costs for patients.^26^ Accordingly, based on practical utility, the approach presented in the current article is expected to offer a more attractive alternative for people willing to assess the intensity of their aging processes.

Epigenetic aging clocks remain the gold standard of aging research due to their high reproducibility and many studies validating their biological relevance.^27^ Although we could not directly compare our aging clock to epigenetic clocks, we identified a statistical association between prediction error and several health conditions, such as hypertension, heart attack, and obesity, as well as a family history of diabetes. These conditions are commonly associated with the aging process and linked to its molecular hallmarks.^28,29^ Specifically, obesity and diabetes represent the faulty nutrient-sensing aspect of aging, while heart conditions are linked to inflammaging, dysregulated cell communication, and mitochondrial dysfunction.^30,31^ These associations indicate that our aging clock is sensitive to these hallmarks despite being based on organismal properties.

Epigenetic aging clocks have been shown to detect the deceleration of aging in response to dietary intervention, supplementation with known geroprotectors, or increased physical activity.^32–36^ We were limited by the retrospective, cross-sectional setting of NHANES and could not assess lifestyle changes’ effects on the pace of aging. However, we expect the presented model to be useful in lonfitudinal settings as well. DASH (Dietary Approaches to Stop Hypertension) or calorie-restrictive diets and exercise affect both blood biometrics and the blood biomarker variables featured in this study’s clock; thus, their effect can translate into a shift in predicted age.^37,38^ Since the reported aging clock is strongly associated with cardiovascular conditions, we believe its predictions may be used to identify people whose heart health requires increased attention. Higher physical activity levels and antihypertensive diets, such as DASH, would be recommended for people with accelerated aging rates. Substance abuse, represented by tobacco-smoking and drinking more than once a week, were the key contributors to aging rates identified in this study. Since we identified no significant association between aging rates and totally abstaining from smoking or drinking, we expect alcohol and tobacco’s effects on the pace of aging to be partly reversible.

The most important feature for biological age prediction was glycated hemoglobin, a blood parameter representing the carbohydrate metabolism that is used in diagnosing diabetes.^39^ Other important features include systolic pressure, waist circumference, and total cholesterol—well-established risk factors for cardiovascular conditions and metabolic syndrome, whose incidence is higher among older adults.^40–42^

These blood biomarkers should, therefore, be monitored regularly in fast agers. Hypertension management is a major focus of healthcare policies in developed countries, such as the US where antihypertensives constitute 10% of total drug expenditure ^43^. According to the PREMIER study concluded in 2003, DASH and lifestyle interventions can reduce systolic blood pressure by 10.5-11.1 mmHg ^44^. The tested lifestyle interventions included maintaining BMI<25 kg/m^2^, lowering sodium intake (e.g. by cutting down on processed foods), performing moderate physical activity for 180 minutes per week, and reducing alcohol consumption. Communicating the benefits of these actions and improving health literacy has been highlighted as one of the key ways to enhance hypersion control ^45^.

Accordingly, we believe the aging clock described in this article may be used as a proxy for cardiovascular and metabolic health, with lower biological age measurements representing lower health risks. When used in clinical practice, this clock can comprehensively measure physical fitness, which can be communicated to patients in order to convey the importance of following prescribed treatments.

## Acknowledgements

Not Applicable

## Ethics Statement

The study was conducted using publicly available data from NHANES. Data collection was observed and approved by the NCHS Ethics Review Board.

## Author contributions

DN — Project administration, Supervision, Resources

FG — Writing – Original Draft Preparation, Supervision, Writing – Review & Editing, Visualization, Methodology, Software

SKG — Investigation, Data Curation, Formal Analysis, Software

## Informed Consent

Not Applicable

